# Single-cell transcriptomics reveals functional insights into a non-model aquatic phytoflagellate and its metabolically linked bacterial community

**DOI:** 10.1101/2023.08.31.555713

**Authors:** Javier Florenza, Aditya Jeevannavar, Anna-Maria Divne, Manu Tamminen, Stefan Bertilsson

**Author notes:** These authors contributed equally to this work (Javier Florenza, Aditya Jeevannavar, Manu Tamminen, Stefan Bertilsson). Department of Organismal Biology, Uppsala University, Uppsala, Sweden (Javier Florenza).

## Abstract

Single-cell transcriptomics is a vital tool for unraveling metabolism and tissue diversity in model organisms. Its potential for elucidating the ecological roles of microeukaryotes, especially non-model ones, remains largely unexplored. This study employed the Smart-seq2 protocol on *Ochromonas triangulata*, a microeukaryote lacking a reference genome, showcasing how transcriptional states align with growth phases. Unexpectedly, a third transcriptional state was identified, across both growth phases. Metabolic mapping revealed a down-regulation trend in pathways associated with ribosome functioning, CO2 fixation, and carbohydrate catabolism from fast to slow growth to the third transcriptional state. Using carry-over rRNA reads, taxonomic identity of *Ochromonas triangulata* was re-confirmed and distinct bacterial communities associated with transcriptional states were identified. This study underscores single-cell transcriptomics as a powerful tool for characterizing metabolic states in microeukaryotes without a reference genome, offering insights into unknown physiological states and individual-level interactions with different bacterial taxa. This approach holds broad applicability for uncovering ecological roles, surpassing alternative methods like metagenomics or metatranscriptomics.

## Introduction

Microbial eukaryotes comprise the vast majority of known eukaryotic lifeforms (Simpson et al. 2017). They showcase a plethora of lifestyles, from free living to strictly parasitic (Walker et al. 2011), photosynthetic, phagotrophic or both (Worden et al. 2015, Mitra et al. 2016). They exhibit a varied catalogue of gene expression quirks, such as mRNA fragmentation, *trans*-splicing or translational slippage (Smith and Keeling 2016). They are ubiquitous across various aquatic and terrestrial environments (Geisen et al. 2015, 2016, Pernice et al. 2016), ranging from freshwater to extreme habitats (Iniesto et al. 2022, Jamy et al. 2022), tropical to polar (Marshall and Laybourn-Parry 2002, Mahé et al. 2017), and are furthermore subject to very strong spatial (Forster et al. 2016) and seasonal constraints (Moreira and López-García 2019, Vass et al. 2020). The unifying feature that groups them together, beyond their eukaryotic characteristics, is that they are single celled and, therefore, microscopic. Their small size, coupled to the difficulty of culturing the overwhelming majority of known single-celled eukaryotic organisms (Del Campo et al. 2014), has technically limited the study of their taxonomy, physiology, and ecology to those species that could be readily cultured or visually recognized under the microscope.

The advent and subsequent development of techniques based on nucleic acid sequencing has circumvented these limitations, greatly expanding our understanding of microeukaryote biology (Goffeau et al. 1996, Venter et al. 2004). During the last two decades, several *en masse* approaches based on molecular characterization (i.e. “omics” methods) have developed into pivotal tools to study the composition and functional potential of microbial communities. Many of these approaches, however, lack single-cell resolution and therefore cannot capture heterogeneity at organismal level. Single-cell methods have emerged to overcome this limitation. The advantages of targeting individual cells are multiple: assays require little starting material, are independent of culture availability, are not limited by high microbial community diversity and provide resolution at organismal level. In the case of single cell mRNA sequencing (scRNA-seq), single cell resolution allows accessing transcriptomic information that would otherwise be too scarce to be recovered from bulk extraction (for example, when the abundance of target organisms is low), and can resolve coexisting, conspecific cell types with disparate gene expression profiles which otherwise would be averaged in an integrated sample.

Despite this potential, studies targeting gene expression in microeukaryotes at single cell levels are scarce, and mostly committed to either yeast model organisms (Nadal-Ribelles et al. 2019, Saint et al. 2019) or pathogenic apicomplexans, such as *Plasmodium* (Poran et al. 2017, Reid et al. 2018, Howick et al. 2019) and *Toxoplasma* (Xue et al. 2020). To date, only two publications exist that offer insight into microeukaryote gene expression beyond either of these lineages: one exploring the feasibility of expression-targeted scRNA-seq in two small flagellates, the haptophyte *Prymnesium parvum* and the dinoflagellate *Karlodinium veneficum* (Liu et al. 2017) and another uncovering successive stages of infection in the haptophyte *Emiliania huxleyi* by its specific giant virus (Ku et al. 2020).

This paucity of studies is partly a consequence of methodological constraints and challenges associated with single cell transcriptomic techniques, such as the need for rapid cell isolation upon sampling, low throughput of isolation strategies compatible with large cells where manual cell isolation is the only feasible option, the low number of mRNA molecules per individual cells or the difficulty of lysing walled or shielded cell types (Liu et al. 2017, Ku and Sebé-Pedrós 2019). Despite these limitations, the *Plasmodium* and *Emiliania* examples showcase scRNA-seq as a feasible approach to metabolically map distinct cell stages within complex microbial populations. However, both examples rely on the availability of either good reference genomes or an integrated reference transcriptome to which scRNA-seq reads can be mapped to build single cell expression profiles. Given the difficulty of culturing most of the enormous diversity of single-celled eukaryotic life forms, such resources are typically not available. In these cases, an *ad hoc*, partial transcriptomic reference can still be built by assembling *de novo* the whole set of mRNA reads from the total pool of captured cells. In this study, we aim to test if such an approach is feasible. For this purpose, we use full-length mRNA sequencing to explore and analyse expression profiles of the single celled chrysophyte *Ochromonas triangulata*, for which no genome reference is available. Cells from two distinct growth phases in culture were FACS-sorted and single-cell transcriptomic libraries were prepared following the Smart-seq2 protocol, which provides the full-length transcript coverage needed for *de novo* assembly of its transcriptome.

## Materials and methods

### Culture conditions

Clonal, unialgal, non-axenic stock cultures of the mixotrophic chrysophyte, *O. triangulata* (∼8 μm in cell diameter) strain RCC21 (Roscoff Culture Collection), were grown in batches of filter-sterilized K/2 medium (Keller et al. 1987, modified by substituting natural seawater for artificial seawater prepared according to Berges et al. 2001). The cultures were serially kept at 18°C under a 12 h photoperiod (120 μmol m^-2^ s^-1^ photon flux measured with a QSL-100 spherical sensor, Biospherical Instruments) by diluting 10 times a fraction of each batch into fresh medium approximately every ten days. Following this routine, representative cells from the fast-growing phase (0.6 doublings per day) were sampled after 2 days from the refreshing date, whereas representatives of the slow-growing phase (0.2 doublings per day) were sampled 11 days after refreshing. *O*. *triangulata* cell abundance prior to sorting was monitored daily using a CytoFLEX flow cytometer (Beckman Coulter). Cell discrimination was based on both side scatter signal (SSC) and chlorophyll *a* autofluorescence detected in the FL3 channel (EX 488, EM 690/50). Cell abundance of the co-cultured, indigenous bacterial community was based on fixed samples (1% formaldehyde final concentration) kept at 4°C upon fixation and analysed with the Cytoflex within 24 h from sampling after staining with SYBR™ Green I (Invitrogen) and jointly detected by SSC and FL1 fluorescence (EX 488, EM 525/40).

### Sorting

Cells from both growth phases were stained with LysoSensor™ Blue DND-167 (Invitrogen), used both as an indicator of cell viability and for presence of food vacuoles. Single cell sorting was performed with a MoFlo Astrios EQ cell sorter (Beckman Coulter) using 355 nm and 640 nm lasers for excitation, 100 μm nozzle, sheath pressure of 25 psi and 0.1 μm sterile filtered 1 x PBS as sheath fluid. Side scatter was used as trigger channel. Sort decisions were based on gates indicating presence of chlorophyll *a* (640-671/30 vs SSC) in combination with LysoSensor detection (355-448/59, Supplementary Figure 2). Singlets were cytometrically selected based on the height-to-area relationship of the pulse (640-671/30 Height-log vs 640-671/30 Area-log). Individual cells were sorted based on the most stringent single cell sort settings available (single mode, 0.5 drop envelope) and deposited into 384-well Eppendorf twin.tec^®^ PCR plates containing 2 μL of lysis buffer (see Library preparation section for composition). The sample plates were divided into four regions of equal size and cells from either growth phase were distributed in alternating regions. This was made to even out technical variation derived from downstream plate processing. A validation plate serving as an initial test of success was sorted using two columns with cells from each of the growth stages. The first two wells in each validation column contained 10 and 5 cells, respectively, to be used as positive controls. The plate holder of the sorter was kept at 4°C at all times. After the sort, the plates were immediately spun down and kept on dry ice until transfer to −80°C for storage. Flow sorting data was interpreted and displayed using the sorter-associated software Summit v 6.3.1.

### Library preparation

All plates were processed according to the Smart-seq2 protocol (Picelli et al. 2014) with slight modifications to comply with small-volume handling robots. The oligo-dT (Smart-dTV30, 5’-Biotin-AAGCAGTGGTATCAACGCAGAGTACT30VN-3’) primer, template switching oligo (TSO, 5’-Biotin-AAGCAGTGGTATCAACGCAGAGTACATrGrG+G-3’) primer, and pre-amplification (IS-PCR, 5’-Biotin-AAGCAGTGGTATCAACGCAGAGT-3’) primer were all modified with a 5’-biotin. This is crucial to increase cDNA yield by avoiding concatenation of TSOs after the first stand reaction (Kapteyn et al. 2010, Turchinovich et al. 2014). The lysis buffer contained a final concentration of 0.2 % Triton -X100 (Sigma), 1 U/μL RNAse inhibitor (cat. no. 2313A, Takara), 2 mM dNTPs (ThermoFisher), Smart-dTV30 primer (IDT) and ERCC RNA Spike-in Mix (cat. no. 4456740, ThermoFisher) diluted 4·10^5^ times. The lysis buffer was cold-dispensed in 2 μL fractions using the MANTIS liquid dispenser (Formulatrix). Plates with sorted cells in lysis buffer were thawed and cDNA generation was conducted in 5 μL reactions containing a final concentration of 1x Superscript II buffer, 5 mM DTT, 1 M MgCl_2_, 1 U/μL RNAse inhibitor (cat. no. 2313A, Takara), 5 U/μL SuperScript™ II Reverse Transcriptase (Invitrogen) and 1 μM TSO (Qiagen). The master mix was dispensed using the MANTIS liquid dispenser followed by mixing for 1 min at 1800 rpm on a plate shaker (Biosan). First strand reaction was run at 42°C for 90 min, followed by 10 cycles of 50°C for 2 min and 42°C for 2 min, with a final 5 min extension at 72°. A 72°C initial denaturation step was omitted as this had shown to have no effect on the results.

Pre-amplification was performed in 12.5 μL final volume with a final concentration of 1x KAPA HiFi HS RM (Roche) and 10 μM IS-PCR primer. Following 3 min at 98°C, amplification was for 24 cycles of 98°C for 20 s, 67°C for 15 s and 72°C for 6 min, with a 6 min final extension at 72°C. Bead cleanup of cDNA was automated using the Biomek NXP liquid handler (Beckman Coulter), AMpure beads and an Alpaqua 384 post magnet with spring cushion technology (Alpaqua). In short, 10 μL beads were added to the 12.5 μL cDNA, the plate was mixed for 1 min at 1800 rpm on a plate shaker and incubated for 8 min before spinning down the liquid and placing the plate on a magnet for 5 min. While on the magnet, the supernatant was removed and 25 μL of freshly prepared 80% EtOH was added and then aspirated after 30 s. The washing was repeated and the plate was left to dry for 2-3 minutes. Elution of cDNA was done by adding 13 μL of water to the wells and the plate was then mixed before 5 min of incubation at RT. Plates were briefly centrifuged and then put on the magnet for 2 min before transfer of 10 μL to a new plate.

Twenty-two single reactions were randomly chosen across the plate for quantification using Qubit HS DNA ready mix, of which 11-15 samples were also run on the Bioanalyzer using the HS DNA chip (Agilent).

### Sequencing

Nextera XT libraries were prepared in 5 μL reactions where all reagent volumes had been scaled down tenfold. The purified cDNA was diluted to approximately 200 pg/μL based on the average cDNA concentration of the 22 randomly chosen reactions. The ATM and TD reagents were pre-mixed and distributed in a 384-well plate before adding cDNA using the Mosquito HV contact dispenser (SPT Labtech). Barcodes of the Illumina Nextera^TM^ DNA UD index set A-D (Illumina) were diluted 1:5 prior to the library amplification. Reactions were run for 12 cycles according to cycling conditions recommended by the manufacturer. For all steps, including manual mixing, a plate shaker run at 1800 rpm and a plate spinner were used. Single reactions from each plate were pooled and purified using 1:0.8 AMPure XP beads. The two pools from each plate were run on separate lanes on a NovaSeq 6000 SP v1.5, PE 1 x 150 bp, including 1% PhiX spike-in. All sequence data generated in this project have been deposited in the European Nucleotide Archive (ENA) at EMBL-EBI and made publicly available under accession number PRJEB60973.

### De novo transcriptome assembly and read mapping

Trinity (Grabherr et al. 2011) was used for *de novo* combined assembly of the *O. triangulata* partial transcriptome as a reference. Reads from all the individual cells were combined for this task. Following assembly, reads from individual cells were aligned to the reference transcriptome using Bowtie (Langmead et al. 2009) and transcript abundance was estimated using RSEM (Li and Dewey 2011).

### Clustering and transcript analysis

Before clustering, data from cells with very low read coverage (fewer than 50 000 reads) were removed. Additionally, contigs that were observed in only a single cell or had over 20 million TPM (Transcripts Per Million) across the population were filtered out to avoid under- and over-representation bias, respectively. The transcript count matrix was dimensionally reduced using *t*-SNE (*t*-distributed Stochastic Neighbourhood Embedding). Further, clustering was performed using the DBSCAN (Density-Based Spatial Clustering of Applications with Noise) algorithm (5 minimum points within ε = 4.5). Transcript accumulation and transcript coverage curves following rarefaction were generated using the iNEXT R package (Hsieh et al. 2016) analogously to procedures used for obtaining curves of species accumulation and species coverage (Chao et al. 2014), using as counterparts transcript identity for species identity and cell identity for sample site.

### Transcriptome annotation

The *de novo* assembled transcriptome was annotated using blastx as implemented in the BLAST+ suite (Camacho et al. 2009). UniProtKB/Swiss-Prot was used as the database for the annotation. MGkit (Rubino et al. 2014), AGAT (v1.1.0, Dainat et al. 2021) and TransDecoder (v5.5.0, Haas 2018) were used to obtain only the annotated parts of the assembled contigs. The longest isoforms were selected using a custom python script. Reads from the individual cells were re-mapped to the annotated transcriptome for further expression analysis.

### Differential expression analysis

Pairwise differential expression analysis was carried out using DESeq2 (Love et al. 2014). The analysis was performed with reads mapped to both the assembled transcriptome and annotated transcriptome separately. Transcripts with a Benjamini-Hochberg adjusted p-value less than 0.05 were considered as differentially expressed.

### Metabolic mapping

The expression matrix of annotated transcripts was mapped to the Kyoto Encyclopedia of Genes and Genomes (KEGG) Pathway database. Pathview (Luo and Brouwer 2013) was used to map mean expression levels of proteins in the three clusters to metabolic maps or pathways obtained from the KEGG Orthology database. To reduce the number of maps to a tractable number, only maps with at least 1 matched protein, 0.25 match ratio (number of matched proteins/ number of proteins in pathway), and 0.1 differential expression ratio (number of differentially expressed proteins/ number of matched proteins) were retained. The remaining pathways were manually inspected and pathways where the only differentially expressed genes were not specific to the pathway were removed.

### 3D structure-based transcriptome annotations

70447 open reading frames, identified using TransDecoder (v5.5.0, Haas 2018), were selected from 150823 transcripts after filtering out transcripts that were present in a single sample or had a cumulative TPM of over 20000000. Of these, very large proteins (>800aa) were excluded, and 3D structures were generated for 66 973 proteins using ESMFold (Lin et al. 2023). After removing structures with low confidence (pLDDT <= 0.5), 54 721 proteins remained. Read mapping was done against the corresponding transcript ORFs using RSEM (Li and Dewey 2011). Structural search was performed using Foldseek (van Kempen et al. 2023) against the Protein Data Bank (PDB, Berman et al. 2000) as well as the AlphaFold structures of the Uniprot/Swiss-Prot database (AFDB, Varadi et al. 2022) with an e-value cut-off of 10. Differential expression analysis using DESeq2 (Love et al. 2014) and metabolic mapping using pathview (Luo and Brouwer 2013) was conducted on the reads mapped to these PDB and AFDB annotated proteins. Protein functions, in the form of Enzyme Commission (EC) numbers and Gene Ontology: Biological Processes (GO:BP) terms, were predicted using a graph convolutional network based tool called DeepFRI (Gligorijević et al. 2021). Gene set enrichment analysis was performed using fgsea (Korotkevich et al. 2021) with the predicted GO:BP terms as the genesets and the list of differentially expressed genes as the pre-ranked list.

### Detection and annotation of rRNA reads

Detection of sequence reads originating from rRNA was carried out with RiboDetector (Deng et al. 2022). For taxonomic classification of putative rRNA sequences, reads of 150 nucleotides or longer and represented at least twice in the whole dataset were annotated using the SINTAX algorithm implementation in VSEARCH (Edgar 2016, Rognes et al. 2016) against the PR2 Reference Sequence Database version 4.14.0 (Guillou et al. 2013) for eukaryote identification and against the SILVA 138 SSU Ref NR 99 (Yilmaz et al. 2014) for prokaryotic community identification. Alpha and beta diversity of the associated prokaryotes was analysed using the *R* packages *mia* and *vegan*. Differential abundance was determined by a consensus of DESeq2, LinDA, Maaslin2, ANCOMBC, and ALDEx2 with an adjusted *P* < 0.05 (Benjamini-Hochberg correction).

## Results

### De novo transcriptome assembly reveals three distinct metabolic states in unicellular eukaryote

*Ochronomas triangulata* cells sampled from two distinct growth stages (Figure 1A) were sorted based on the joint signal of chlorophyll *a* and LysoSensor Blue (Figure 1B). 744 single-cell mRNA libraries were prepared and sequenced yielding a median of 1.4M reads per cell. *De novo* assembly of reads pooled across all cells produced a transcriptome of ∼166000 contigs (median length 432 bases), which reduced to 60 307 (median length 835 bases) after removing contigs that were not observed in at least 2 cells (Supplementary figure 3). t-SNE dimensionality reduction and DBSCAN clustering of expression profiles correctly identified 639 cells with their sorting-based affiliation (Figure 1C). Unexpectedly, a third, as yet uncharacterised, cluster of cells originating from both the fast (37 cells) and slow (38 cells) sort groups were observed and will henceforth be referred to as the uncharacterised cluster/cells.

**Figure 1:**
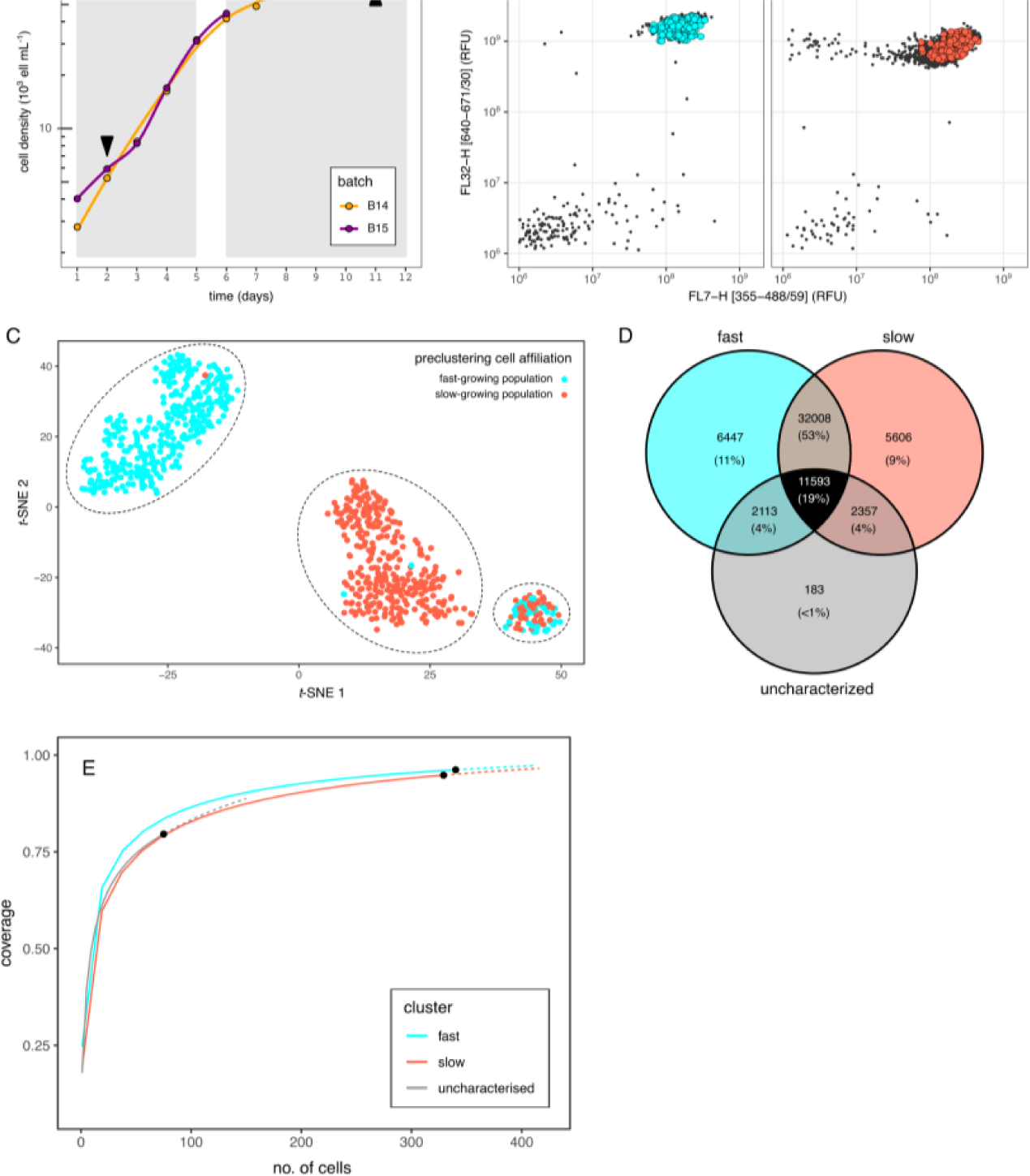
Sampling, sorting, and clustering *Ochromonas triangulata*. (A) Growth curves of source cultures for cells during exponential growth stage (fast growth) and linear growth stage (slow growth). Batch labels refer to the source culture for cells in slow (B14) and fast growth (B15). Arrowheads indicate the sampling points for each stage. (B) Flow cytograms showing placement of FACS-sorted *O. triangulata* cells (coloured dots) for each growth stage. (C) *t*-SNE visualization of expression clusters. Each dot represents one cell color-coded based on its sampling origin – either the fast-growing (cyan) or slow-growing (red) cell consortia. Clusters identified by DBSCAN are enclosed by a dotted line. (D) Venn diagram showing the number of shared and exclusive transcript sets among postclustering groups. (E) Partial transcriptome coverage plot for each postclustering group. The black dots indicate sample size, the solid lines follow sample rarefaction, and the dotted lines represent an extrapolation based on the rarefaction model for each group.

In total, 19% of the contigs were present across all three expression clusters and 53% were shared by only the fast- and slow-growing clusters, while less than 1% were unique to the uncharacterised cluster (Figure 1D). However, all expression clusters showed coverage of cluster-specific transcriptomes above 75%, with the fast- and slow-growing clusters, having more representatives, very close to full transcriptome completeness (Figure 1E)

### Core metabolic activity occurs in three distinct levels in the three clusters

Only 17883 transcripts out of the 60307 contigs could be annotated using sequence homology against the UniProt database, corroborating the underrepresentation of annotated protist sequences in public databases (Keeling et al. 2014). Pairwise comparisons among the three distinct cell clusters revealed 538 differentially expressed annotated genes (DEGs, Figure 2). Mapping these DEGs against KEGG Orthology pathways identified four main pathway families: transcription and translation processes, vesicle maintenance and membrane trafficking, photosynthetic activity, and carbohydrate metabolism (Figure 3). DEGs in the KEGG pathways detected from any pairwise comparison against the uncharacterised group were in all instances and without exception, downregulated in the latter.

**Figure 2:**
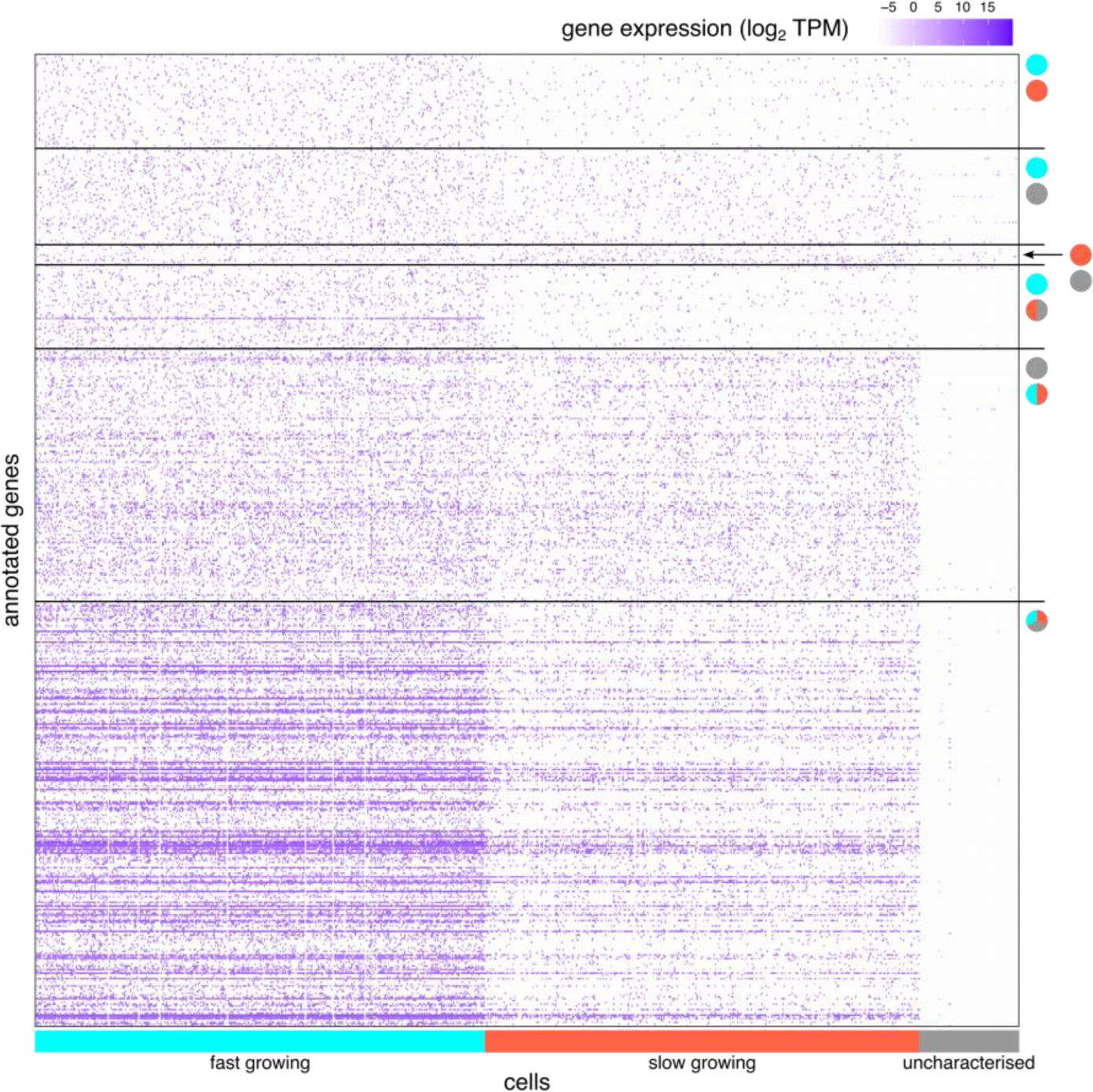
Heatmap of annotated genes that were differentially expressed between different *Ochromonas triangulata* growth stages. Coloured rectangles in the horizontal axis indicate cell affiliation to clustering group. Circle pairs indicate pairwise comparisons between stages. Those cases where gene expression in one group is significantly different to that of the other two groups, the joint is represented by a split circle in which the colour of each half encodes the identity of each member. A three coloured circle represents cases in which gene expression differs significantly for any possible pairwise comparison. The colours in circles and rectangles correspond to fast-growing (cyan), slow-growing (red) and uncharacterised (grey) stages.

**Figure 3:**
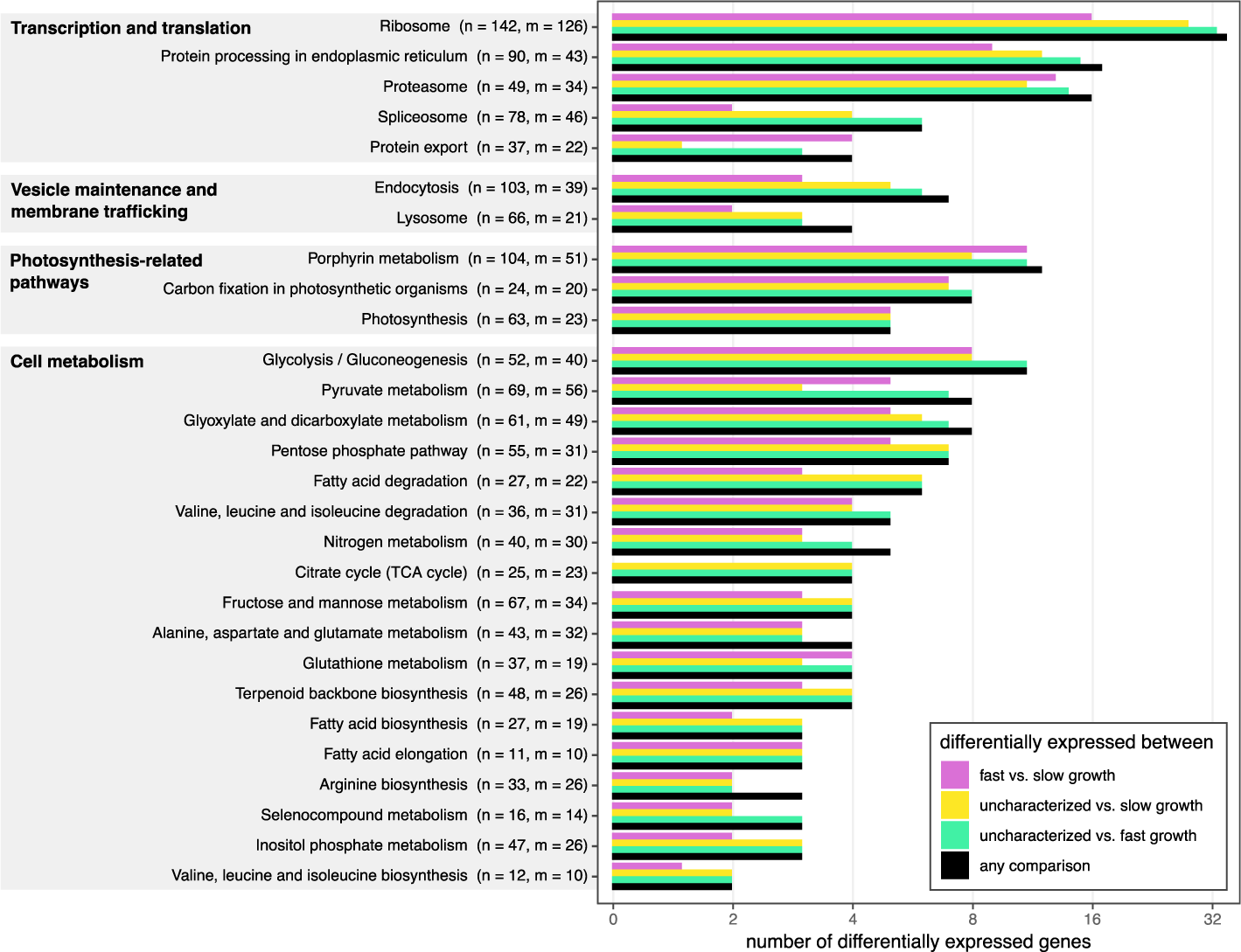
Summary of KEGG pathways associated to multiple differentially expressed genes. For each pathway label, *n* indicates the total number of genes that belong to the pathway and *m* the number of genes for which expression was recovered in our transcriptomes.

Three pathways related to transcription and translation (ribosome, protein processing and proteasome) featured the highest number of differentially expressed genes. In addition, two other key components in the progression from transcript to functional protein (spliceosome, protein export) also had a small number of differentially expressed genes. Downregulation of genes involved in such processes were prevalent in the uncharacterised cells (28 and 33 genes when comparing it to the slow-growing or fast-growing cells, respectively). This agrees with the expression level pattern across the groups observed in Figure 2 and suggests reduced transcriptional activity in the uncharacterised group when compared to the other two.

The second pathway family (endocytosis and lysosome pathways) provides insights into phagocytotic behaviour, revealing joint downregulation of clathrin and AP-2 (both essential coating components of vesicles resulting from endocytosis) in the uncharacterised group as well as high expression of lysosomal proteases (homologs of tripeptidyl-peptidase 1 and cathepsin A, B, D, F and X) in both fast and slow-growing expression clusters in parallel with negligible expression of these proteases in the uncharacterised group. These observations, although limited, provide evidence of active lysosomal digestion in the growing cells, which in contrast seems to be absent from the uncharacterised cells.

Finally, the third and fourth pathway families, represented by the pathways associated to photosynthesis and carbohydrate metabolism, harbour the highest number of DEGs featured in the whole set of KEGG and provide the clearest picture associated to physiological differences between cell stages (Figure 4). In all cases, DEGs in this category showed high expression in the fast-growing cells, and low in the uncharacterised cells, while expression was either intermediate or comparable to the fast-growing in the slow-growing cells.

**Figure 4:**
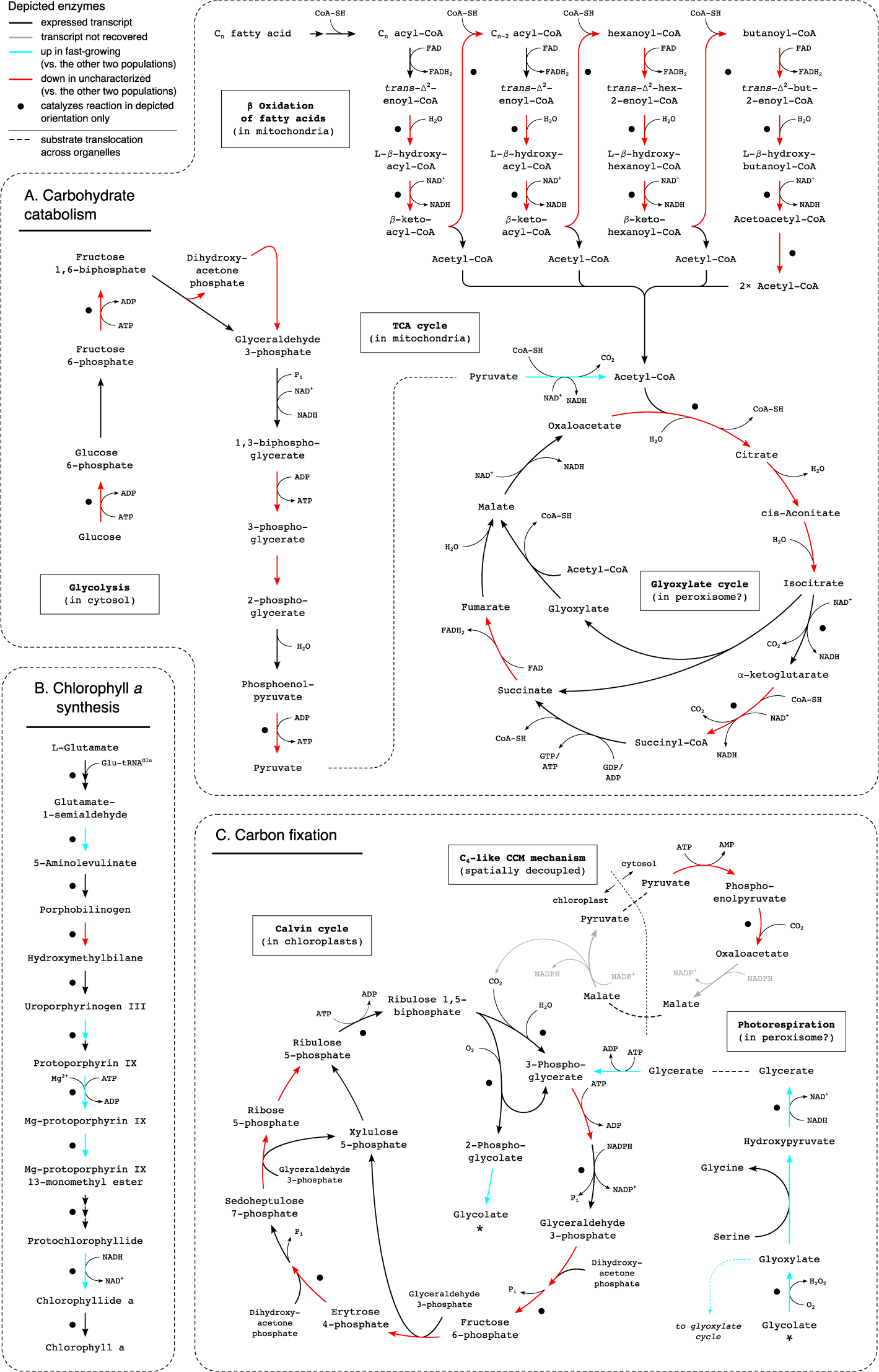
Differential expression in key metabolic pathways involving (A) carbohydrate catabolism (glycolysis, tricarboxylic acid cycle, and β oxidation of fatty acids), (B) chlorophyll *a* synthesis, and (C) carbon fixation (calvin cycle, C_4_-like CCM mechanism, and photorespiration).Coloured arrows represent enzymes that catalyse the depicted reactions. Black dots indicate enzymes that only catalyse the reactions in the depicted direction. Dashed lines indicate substrate translocation across organelle boundaries.

Widespread down-regulation of carbohydrate catabolism is a pronounced feature of the uncharacterised cells, apparent through the downregulation of key enzymes of the glycolysis pathway (phosphofructokinase-1 and pyruvate kinase), the β oxidation of fatty acids (enoyl-Coa hydratase, β-hydroxyacyl-CoA dehydrogenase, and acyl-Coa acetyltransferase) and the tricarboxylic acid (TCA) cycle (citrate synthase) (Figure 4A). Expression of all enzymes involved in β oxidation and most enzymes in the TCA cycle is recovered in all three clusters. Low expression of genes mediating such pathways imply low levels of energy production for cell maintenance in the uncharacterised cluster compared to the other two.

In contrast, observable in the fourth KEGG family pathway with highest number of DEGs (porphyrin metabolism, Figure 3), exacerbation of chlorophyll *a* synthesis manifested in the fast-growing stage (Figure 4B). It involves the upregulation of 5 enzymes in the pathway, including magnesium chelatase, which catalyses the first committed step of chlorophyll *a* production, and protochlorophyllide reductase, which generates chlorophyllide *a*, the immediate precursor of chlorophyll *a*. A putative increased chlorophyll *a* production in the fast-growing stage is in accordance with an enhanced chlorophyll fluorescence signal in the sorting cytograms (Figure 1B).

Finally, downregulated key enzymes of the Calvin cycle in the uncharacterised cells (fructose 1,6-biphosphatase and fructose-bisphosphate aldolase, both catalysing unidirectional reactions, Figure 4C) indicates little or no carbon fixation in this group. At the same time, upregulation of photorespiration enzymes in the fast-growing stage suggests high CO_2_ fixation activity associated to these cells, since this is a compensatory mechanism to counteract the efficiency loss in carbon fixation due to the oxidase activity of rubisco. Photorespiration is expected to occur in peroxisomes (Sandalio and Romero-Puertas 2015), which are not characterized in chrysophytes but are reported in other stramenopiles both based on microscopic and genomic evidence (Gabaldón 2010, Mix et al. 2018). This, in combination with increased chlorophyll *a* production, points towards growth conditions mostly supported by photosynthesis.

Among DEGs not associated to a relevant KEGG Pathway, only three show significant upregulation in the uncharacterised cluster. Their UniProt-based annotation corresponds to prokaryotic entries in the database, although the three genes have also eukaryotic homologs. Additionally, many genes unique to this uncharacterised group (42 of the 183 contigs) all had prokaryotic hits as basal annotation, 22 of which lack any eukaryotic homolog.

### Novel structural homology analysis pipeline adds annotation and upholds sequence homology-based inferences

ESMFold successfully predicted the 3D structures for 54725 proteins (Figure 8E, Supplementary tables 1, 2), including 32858 proteins with high confidence (0.7 < pLDDT <= 0.9) and 8 328 proteins with very high confidence (pLDDT > 0.9). Cells in fast-growing, slow-growing, and uncharacterised clusters featured a median of 1094, 878, and 624 expressed proteins, respectively (Figure 5A). Over 80% of the proteins expressed were common to both fast- and slow-growing clusters while only 0.22% of the predicted proteins were uniquely expressed in the uncharacterised cluster (Figure 5C). The distribution of predicted proteins expressed in the three clusters was similar to the distribution based on sequence homology. A median of 164184, 122519, and 75196 RNA reads mapped to the fast-growing, slow-growing, and uncharacterised cells, based on the predicted proteins’ transcripts, respectively. Overall, these comprised 9.31% of all reads sequenced which is similar to the 10.5% reads mapping to the transcripts annotated by sequence similarity analyses (Figure 5B, Supplementary figure 5).

**Figure 5:**
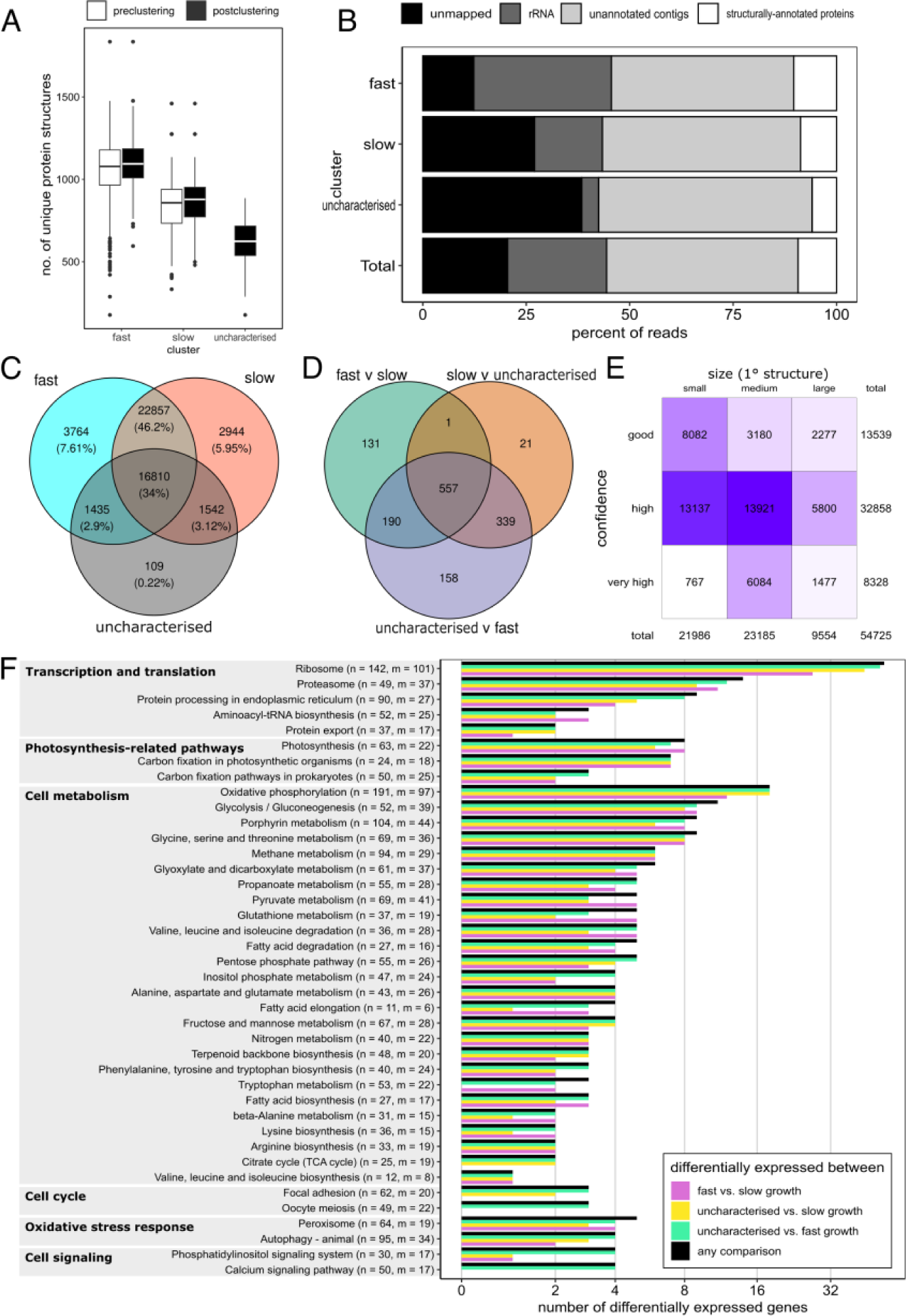
Protein tertiary structure-based analyses. (A) Distributions of protein richness per cell. (B) Relative proportion of total reads that either didn’t map to the assembled transcriptome, were assigned as ribosomal, or mapped to unannotated contigs or structurally-annotated proteins (C) Venn diagram showing the number of shared and exclusive protein sets among postclustering groups (D) Venn diagram showing the number of shared and exclusive protein sets that were differentially expressed among the postclustering groups (E) Distribution of the predicted protein structures on protein sequence length (no. of amino acids: small (0, 200], medium (200, 400], large (400, 800]) and structure confidence (pLDDT scores: good (0.5, 0.7], high (0.7, 0.9], very high (0.9, 1]) axes. (F) Summary of KEGG pathways associated to multiple differentially expressed genes. (n = no. of proteins in pathway, m = no. of those proteins recovered). Functions and pathways also identified by sequence homology (fig. 3) include transcription and translation (ribosome, protein processing in endoplasmic reticulum, and proteasome), photosynthesis (porphyrin metabolism, photosynthesis, and carbon fixation in photosynthetic organisms), and cell metabolism (glycolysis, pyruvate metabolism, citrate cycle, fatty acid biosynthesis and elongation etc.) associated pathway groups.

Unlike sequence similarity-based annotation, where only 29.6% of the transcripts were successfully annotated, over 99.9% of the proteins expressed and folded were successfully annotated based on structural homology. 1397 of these proteins were significantly differentially expressed between the comparisons, with 557 proteins significant in all three comparisons, with high expression in fast-growing cells, intermediate expression in slow-growing cells, and low expression in uncharacterised cells (Figure 5D, Supplementary figure 7). Additionally, there were 339 proteins that had comparable expression between fast- and slow-growing cells, while having low expression in uncharacterised cells.

The KEGG pathways associated to these differentially expressed proteins highly overlapped the set obtained from metabolic mapping following the sequence similarity-based analysis (Figures 5F, 2), representing transcription and translation, photosynthesis, and cell metabolism associated pathway groups. Additionally, pathways associated with cell cycle (focal adhesion and oocyte meiosis) and oxidative stress (autophagy and peroxisome) were identified. This was further evident from gene set enrichment where a total of 100, 91, and 69 GO: BP gene sets were significantly enriched for in the fast- vs. slow-growing, uncharacterised vs. fast-growing, and uncharacterised vs. slow-growing comparisons including GO terms for cellular response to Oxidative stress (GO:0034599), Oxygen radical (GO:0071450), Reactive oxygen species (GO:0034614), Stress (GO:0033554), as well as DNA repair (GO:0006281). Some pathways’ enrichment suggested bacterial symbiosis, for instance Interspecies interaction (GO:0044419) and Symbiotic interaction (GO:0044403) and potentially bacterial invasion, for instance Interaction with host (GO:0051701), Entry into host (GO:0044409), and Movement in host (GO:0052126, Supplementary tables 3, 4, 5).

### rRNA read carry-over enable identification of microeukaryote **and** distinct associated prokaryotes

Overall, 241.7 M reads were predicted to be of rRNA origin across all samples (24% of the total), although rRNA read coverage was uneven across cell types, constituting 33%, 16% and 4% of the fast-growing, slow-growing, and uncharacterised cells, respectively (Supplementary Figure 5). After discarding singletons and reads shorter than 150 nucleotides, the resulting set was successfully dereplicated into 4.5 million unique sequences. Of these, one third (33%) could be taxonomically classified with confidence (>0.8 bootstrap support). Annotated read recovery across samples was sufficient (median 106 688 sequences per sample) to correctly annotate 653 cells (88%) as *Ochromonas triangulata* (median 23 811 per sample). 16 of the remaining 91 cells were excluded from all transcriptomic analysis due to a failure to map at least 50 000 reads to the assembled transcriptome. The remaining 75 cells belonged, without exception, to the uncharacterised cluster.

Median rRNA abundance in the uncharacterised cluster was low (approx. 43 000 reads per cell). A broadly reduced expression landscape, which also includes lower demand for rRNA (as suggested by generalized downregulation of ribosomal proteins, Figure 3) might explain insufficient rRNA coverage for unambiguous annotation of cells characterized by low transcriptional activity.

The 16S rRNA read recovery across the samples was relatively low (median 2 807 reads per sample) but saturated (Supplementary Figure 6) and sufficient to characterize the prokaryotic community composition associated with individual *Ochromonas triangulata* cells.

The prokaryotic communities of the fast-growing and uncharacterised cells were more internally uniform (Figures 6A and 6B; median Bray-Curtis dissimilarities 0.1745 and 0.1498 within the groups, respectively) compared to the slow-growing cells (median Bray-Curtis dissimilarity 0.2891 within the group; Figure 6B). The prokaryotic communities were significantly different between the fast- and slow-growing (PERMANOVA *R^2^*=0.4521 *P*=0.001; median Bray-Curtis dissimilarity 0.4340), fast-growing and uncharacterised (PERMANOVA *R^2^*=0.7786 *P*=0.001; median Bray-Curtis dissimilarity 0.797) and slow-growing and uncharacterised cells (PERMANOVA *R^2^*=0.3127 *P*=0.001; median Bray-Curtis dissimilarity 0.4338). In contrast, bacterial alpha diversity (richness) was not significantly different between any of the three groups (Supplementary Figure 6C).

**Figure 6:**
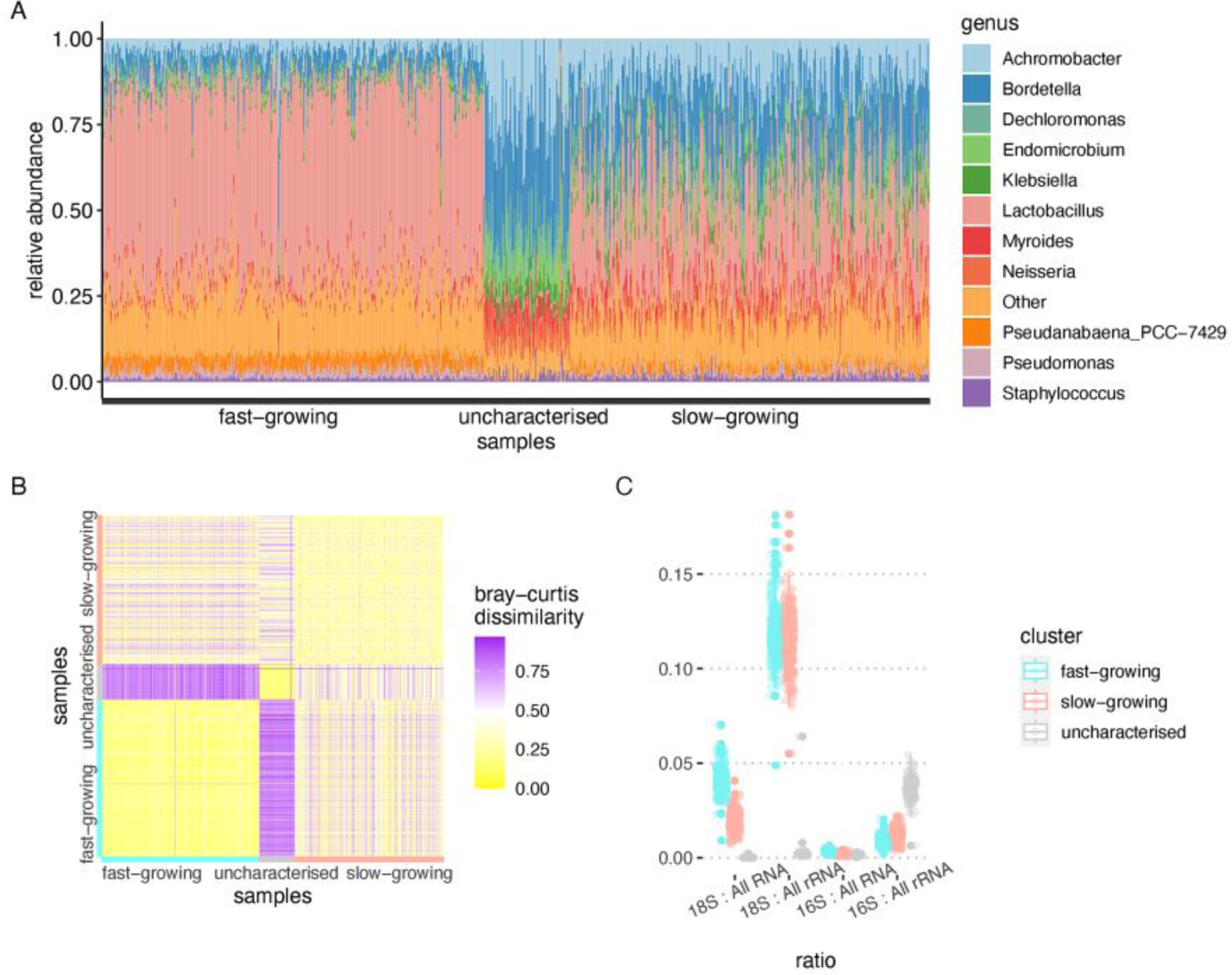
The prokaryota associated with the *O. triangulata* cultures. (A) Relative abundance of the eleven most abundant prokaryotic genera associated with the different expression clusters. (B) Bray-Curtis dissimilarity of each sample’ associated prokaryota with the rest of the samples. (C) Ratios of 18S and 16S rRNA to the rest of the RNA or rRNA obtained from the samples.

The proportion of 16S rRNA to total rRNA was significantly higher in the uncharacterised cell cluster compared to the fast- and slow-growing groups (Wilcoxon *P* < 2.2·10^-16^ and *P* < 0.009, respectively) whereas the proportion of 16S rRNA to total RNA was not significantly different between any of the groups (Figure 6C). This is in contrast with the proportion of the 18S rRNA to all rRNA and all RNA reads, where the 18S rRNA proportion for the uncharacterised group was significantly lower than the others (Wilcoxon test *P* < 2.2·10^-16^ and *P* < 2.2·10^-16^, respectively). The differentially abundant families (Supplementary Figure 6C) between the cell clusters include *Flavobacteriaceae*, *Endomicrobiaceae*, *Alcaligenaceae*, *Burkholderiaceae*, *Neisseriaceae*, *Rhodocyclaceae*, *Enterobacteriaceae* (lowest abundance in the fast-growing population, highest abundance in the uncharacterised population) and *Lactobacillaceae*, *Streptococcaceae*, *Pseudanabaenaceae*, *Rhizobiaceae*, *Anaplasmataceae*, *Pseudomonadaceae* and *Staphylococcaceae* (lowest abundance in the uncharacterised population, highest abundance in the fast-growing population).

## Discussion

Single cell transcriptomic profiling of the microbial eukaryote *O. triangulata* was technically feasible even without any reference genome. The SmartSeq2 protocol, used to generate the mRNA libraries of FACS sorted single cells of *O. triangulata*, provided sufficient output while preserving the full-length transcript information necessary to generate a working transcriptome assembled *ad hoc*. Using this transcriptome as a basis for the analysis of gene expression, we were able to discern three distinct cell clusters of *O. triangulata* originating from two contrasting growth stages, therefore uncovering a third uncharacterised group of cells that was not expected *a priori*.

While we found no technical limitations in deploying scRNA-seq, annotation issues are still pervasive. In our dataset, two thirds of the comprehensive transcriptome remained unannotated (Suppl. Figure 4). Consequently, of all the data generated in our experiment, only approximately 11% of the reads contributed to our understanding of mRNA expression that could be associated with putative biological functions (Suppl. Figure 5). On the other hand, the dataset is characterized by a large proportion of dark expression (i.e. reads that mapped to transcripts devoid of known function, 45% in our dataset), and therefore much biological significance remains unavailable due to weaknesses of annotation. Unavoidably, organisms that are poorly represented in annotation databases (as is the case for most non-model microeukaryotes) will be subject to such constraints.

We explored predictive structural annotation to mitigate this phenomenon. We obtained tertiary structures for ∼55 000 proteins and successfully annotated over 99% of them. Read mapping against these proteins’ transcripts however remained on par, at ∼10%, with mapping against ∼20 000 sequence-annotated transcripts. Such a persistent dark expression after structural analysis could indicate that many contigs and transcripts from the *de novo* assembly may not encode proteins at all.

The problems associated to dark expression are well exemplified by the uncharacterised cell cluster. This cell group had fewer well supported unique transcripts which accounted for a good representation of the expression landscape of the group (covering approx. 80% of its full transcriptome). In turn, most of these transcripts were shared with one or both of the other two groups, while only 1% were unique to this group. The question of what this seemingly uncharacterised cluster represents remain enigmatic. We speculate that these cells might be under colonization by a bacterial community, possibly involving pathogens or parasites. This scenario would be consistent with the prokaryotic nature of at least a portion of the mRNA transcripts unique to the uncharacterised group. This possibility would be compatible with the very erratic recovery associated with these transcripts, since the recovery of non-polyadenylated prokaryotic mRNA, likely caused by random mRNA capture in a similar mechanism driving rRNA leakage, would be expectedly inconsistent. In addition, the relatively high proportion of transcripts that are shared between the uncharacterised cluster and either of the other two groups (slightly above 2 000 transcripts in both cases, Figure 3C) agrees with the hypothesis that the uncharacterised cluster is the result of bacteria invading cells that were originally from the other two groups.

Despite the dark expression, differential expression of genes involved in quasi-universal cellular pathways, and therefore well represented in annotation databases, provided insight into the functional state of *O. triangulata* beyond the uncharacterised group. For example, enhanced photosynthetic activity in combination with seemingly increased CO_2_ fixation were characteristic of cells in fast exponential growth. This, together with a concurrent peak in bacterial abundance during this phase seemed to indicate that a boost in *O. triangulata* growth could be enabled by its heterotrophic activity. When prey declined into a stable, potentially predation-resistant community, *O. triangulata* growth transitioned into a slower growth rate driven mostly by photosynthesis. This would be in line with what has been shown before for *Ochromonas* isolate CCMP1393 (Lie et al. 2018, Wilken et al. 2020), and would delineate *O. triangulata* as a mainly phototrophic constitutive mixotroph that could benefit from prey uptake to accelerate growth (group C mixotroph *sensu* Jones 1997).

Results from the structure-based analysis corroborated those from the sequence-based analysis in terms of read mapping, transcript diversity, and differential expression between clusters, as well as metabolic mapping. Additionally, improved structure and structure-function annotation, based on predicted protein structures, as compared to sequence-based annotation, revealed enrichment in pathways associated with oxidative stress as well as interspecies interaction in the uncharacterised expression cluster, providing further evidence that this uncharacterised population could be composed of cells that are close to dormancy or have been invaded by bacteria. Beyond protein expression, another additional source of useful information that was featured in our dataset came from rRNA annotation. Although Smart-Seq2 targets specifically mRNA, the cellular amount of rRNA in a eukaryotic cell is vast enough to allow considerable leakage of rRNA reads into the dataset. This phenomenon, typically regarded as a nuisance, can be used to the researcher’s advantage when taxonomic affiliation of a cell is unknown, as would be the case from a natural sample. In our dataset, about one quarter of the total reads corresponded to rRNA, although it was not equally distributed across cell clusters. This can become limiting when read coverage per cell is relatively low, since we needed >100 000 reads to correctly annotate *O. triangulata* cells based on direct annotation of rRNA reads. The reason for this might be that the taxonomical information conveyed by these sequences is generally poor because they can originate anywhere in the rRNA operon, much of which is phylogenetically ambiguous. In the case of the uncharacterised group, where only 4% of the reads overall originated from rRNA, annotation becomes misleading. The potential prokaryotic nature of many reads associated to the uncharacterised cell group could also explain the difficulty in proper taxonomic annotation for members of this group. Nevertheless, when read support is high, cells could be taxonomically annotated with confidence.

The carry-over of 16S rRNA reads in the sequencing data permits profiling the bacterial communities associated with individual *O. triangulata* cells. Surprisingly, the community compositions were consistent within the clusters but variable between them despite the similarity of the co-cultured bacterial communities (Fig. 6B). The internally most consistent bacterial communities were amidst the fast-growing and uncharacterised cell groups whereas more variability was observed within the slow-growing group. Interestingly, the bacterial community composition of the members of the uncharacterised expression cluster did not depend on their origin in the fast- or slow-growing cells. Also, the slight bimodality in the alpha diversity of the uncharacterised cells had no connection to the growth stage.

We speculate that the differences in the bacterial communities between the groups could be explained by grazing behaviour of the *O. triangulata* cells. Initially, low abundance of *O. triangulata* cells would allow the bacteria to grow fast under little grazing pressure. With time and consequently increasing numbers of *O. triangulata*, we speculate that the peak density of the bacterial community is consumed and the community transitions to a grazing-resistant steady state (Supplementary Figure 1) forcing *O. triangulata* to progress from a fast-growing to slow-growing state. The increased grazing pressure due to higher numbers of *O. triangulata* could selectively affect the community structure of the co-cultured bacteria, which in turn could result in lower stability as well as a shift from the bacterial community associated with the fast-growing population. Unfortunately, the current study lacks community profiling of the co-cultured bacterial cells and therefore we are not able to confirm the similarity of the individual *O. triangulata* bacterial communities to the bulk bacterial community structure of the fast- and slow-growing culture stages.

The specific bacterial community composition within the uncharacterised cell cluster is consistent with the scenario described above, in which bacteria might be colonizing compromised *O. triangulata* cells. This is supported, in addition to the lower abundance of total RNA reads associated to this cluster, by its elevated cell-associated bacterial load, observed as a significantly higher proportion of prokaryotic rRNA in cells from this cluster compared to the other two groups. As the cells in the uncharacterised cluster do not seem to be actively feeding according to the downregulation of key lysosome-related enzymes, colonization by invasive bacteria seems a plausible explanation of the higher prokaryotic rRNA. Finally, the consistent bacterial community composition associated to these cells could be explained by very specific bacterial community members taking part in the invasion.

Overall, our results show that scRNA-seq is technically apt to deal with small microeukaryotes, even those lacking a reference genome. While potential for detailed mechanistic understanding of functional states is limited for organisms that are poorly represented in public databases, such as microeukaryotes, potential for discovery of unexpected and diverse expression consortia remains a key asset of scRNA-seq. Taken together, the combination of 18S rRNA annotation with expression-driven cell clustering and structural homology-driven protein annotation holds promise as a powerful tool to characterize community structure and function from natural samples beyond metabarcoding.

## Acknowledgements

This work was partly financed by the EU’s Horizon 2020 research and innovation programme under the Marie Skłodowska-Curie ITN project SINGEK (grant agreement no. H2020-MSCA-ITN-2015-675752), the Swedish Research Council (grant 2017-04422), Formas (grant 2019-02366) and the Academy of Finland (grant 336475). The authors would like to thank Ian Probert (Roscoff Culture Collection) for providing the strain used in this study. The authors would like to acknowledge support from the Genomics infrastructure services at Science for Life Laboratory. Single Cell Sorting and library preparation was performed at the Microbial Single Cell Genomics Facility (MSCG). Sequencing was performed by the SNP&SEQ Technology Platform. The facilities are part of the Uppsala node at the National Genomics Infrastructure (NGI) Sweden and Science for Life Laboratory. The SNP&SEQ Platform is also supported by the Swedish Research Council and the Knut and Alice Wallenberg Foundation. The Eukaryotic Single Cell Genomics unit at the Stockholm node, Science for Life Laboratory, provided instrumental advice when adapting the SmartSeq2 protocol to low volume reactions. The authors also wish to acknowledge CSC – IT Center for Science, Finland, for computational resources.

## Data Availability Statement

All sequence data generated in this project have been deposited in the European Nucleotide Archive (ENA) at EMBL-EBI and made publicly available under accession number PRJEB60973. They are available at the following URL: https://www.ebi.ac.uk/ena/browser/view/PRJEB60973

## References

Berges, J.A., Franklin, D.J. and Harrison, P.J. (2001) ‘Evolution of an artificial seawater medium: Improvements in enriched seawater, artificial water over the last two decades’, Journal of Phycology, 37(6), pp. 1138–1145.

Berman, H.M. et al. (2000) ‘The Protein Data Bank’, Nucleic Acids Research, 28(1), pp. 235–242.

Camacho, C. et al. (2009) ‘BLAST+: Architecture and applications’, BMC Bioinformatics, pp. 1–9.

Del Campo, J. et al. (2014) ‘The others: our biased perspective of eukaryotic genomes’, Trends in Ecology and Evolution, 29(5), pp. 252–259.

Dainat, J., Hereñú, D. and Pucholt, P. (2021) ‘AGAT: Another Gff Analysis Toolkit to handle annotations in any GTF/GFF format’.

Deng, Z.L., Münch, P.C., Mreches, R. and McHardy, A.C. (2022) ‘Rapid and accurate identification of ribosomal RNA sequences via deep learning’, Nucleic Acids Research, 50(10), p. E60.

Edgar, R. (2016) ‘SINTAX: a simple non-Bayesian taxonomy classifier for 16S and ITS sequences’, bioRxiv [Preprint], (074161).

Forster, D. et al. (2016) ‘Benthic protists: the under-charted majority Dominik’, FEMS Microbiology Ecology, 92, pp. 1–11.

Geisen, S. et al. (2015) ‘Metatranscriptomic census of active protists in soils’, ISME Journal, 9(10), pp. 2178–2190.

Geisen, S. et al. (2016) ‘The soil food web revisited: Diverse and widespread mycophagous soil protists’, Soil Biology and Biochemistry, 94, pp. 10–18.

Gligorijević, V. et al. (2021) ‘Structure-based protein function prediction using graph convolutional networks’, Nature Communications, 12(1).

Goffeau, A. et al. (1996) ‘Life with 6000 Genes’, Science, 274, pp. 546–567.

Grabherr, M.G. et al. (2011) ‘Full-length transcriptome assembly from RNA-Seq data without a reference genome’, Nature Biotechnology, 29(7), pp. 644–652.

Guillou, L. et al. (2013) ‘The Protist Ribosomal Reference database (PR2): A catalog of unicellular eukaryote Small Sub-Unit rRNA sequences with curated taxonomy’, Nucleic Acids Research, 41(D1), pp. 597–604.

Haas, B.J. (2018) ‘TransDecoder v5.5.0’. Available at: https://github.com/TransDecoder/TransDecoder.

Howick, V.M. et al. (2019) ‘The malaria cell atlas: Single parasite transcriptomes across the complete *Plasmodium* life cycle’, Science, 365(6455).

Hsieh, T.C., Ma, K.H. and Chao, A. (2016) ‘iNEXT: an R package for rarefaction and extrapolation of species diversity (Hill numbers)’, Methods in Ecology and Evolution, 7(12), pp. 1451–1456.

Iniesto, M. et al. (2022) ‘Planktonic microbial communities from microbialite-bearing lakes sampled along a salinity-alkalinity gradient’, Limnology and Oceanography, 67(12), pp. 2718–2733.

Jamy, M. et al. (2022) ‘Global patterns and rates of habitat transitions across the eukaryotic tree of life’, Nature Ecology and Evolution, 6(October), pp. 1458–1470.

Jones, H.L.J. (1997) ‘A classification of mixotrophic protists based on their behaviour’, Freshwater Biology, 37(1), pp. 35–43.

Kapteyn, J., He, R., McDowell, E.T. and Gang, D.R. (2010) ‘Incorporation of non-natural nucleotides into template-switching oligonucleotides reduces background and improves cDNA synthesis from very small RNA samples’, BMC Genomics, 11(1).

Keeling, P.J. et al. (2014) ‘The Marine Microbial Eukaryote Transcriptome Sequencing Project (MMETSP): Illuminating the Functional Diversity of Eukaryotic Life in the Oceans through Transcriptome Sequencing’, PLoS Biology, 12(6).

Keller, M.D., Selvin, R.C., Claus, W. and Guillard, R.R.L. (1987) ‘Media for the culture of oceanic ultraphytoplankton’, Journal of Phycology, 23(4), pp. 633–638.

van Kempen, M. et al. (2023) ‘Fast and accurate protein structure search with Foldseek’, Nature Biotechnology [Preprint].

Korotkevich, G. et al. (2021) ‘Fast gene set enrichment analysis’, bioRxiv [Preprint].

Ku, C. et al. (2020) ‘A single-cell view on alga-virus interactions reveals sequential transcriptional programs and infection states’, Science Advances, 6(21).

Ku, C. and Sebé-Pedrós, A. (2019) ‘Using single-cell transcriptomics to understand functional states and interactions in microbial eukaryotes’, Philosophical Transactions of the Royal Society B: Biological Sciences, 374(1786).

Langmead, B., Trapnell, C., Pop, M. and Salzberg, S.L. (2009) ‘Ultrafast and memory-efficient alignment of short DNA sequences to the human genome’, Genome biology, 10(3), p. R25.

Li, B. and Dewey, C.N. (2011) ‘RSEM: accurate transcript quantification from RNA-Seq data with or without a reference genome’, BMC Bioinformatics, 12(323), pp. 1– 16.

Lin, Z. et al. (2023) ‘Evolutionary-scale prediction of atomic-level protein structure with a language model’, Science, 379(6637), pp. 1123–1130.

Liu, Z. et al. (2017) ‘Single-cell transcriptomics of small microbial eukaryotes: Limitations and potential’, ISME Journal, 11(5), pp. 1282–1285.

Love, M.I., Huber, W. and Anders, S. (2014) ‘Moderated estimation of fold change and dispersion for RNA-seq data with DESeq2’, Genome Biology, 15(12), pp. 1–21.

Luo, W. and Brouwer, C. (2013) ‘Pathview: an R/Bioconductor package for pathway-based data integration and visualization’, Bioinformatics, 29(14), pp. 1830–1831.

Mahé, F. et al. (2017) ‘Parasites dominate hyperdiverse soil protist communities in Neotropical rainforests’, Nature Ecology and Evolution, 1(4).

Marshall, W. and Laybourn-Parry, J. (2002) ‘The balance between photosynthesis and grazing in Antarctic mixotrophic cryptophytes during summer’, Freshwater Biology, 47(11), pp. 2060–2070.

Mitra, A. et al. (2016) ‘Defining Planktonic Protist Functional Groups on Mechanisms for Energy and Nutrient Acquisition: Incorporation of Diverse Mixotrophic Strategies’, Protist, 167(2), pp. 106–120.

Moreira, D. and López-García, P. (2019) ‘Time series are critical to understand microbial plankton diversity and ecology’, Molecular Ecology, 28(5), pp. 920–922.

Nadal-Ribelles, M. et al. (2019) ‘Sensitive high-throughput single-cell RNA-seq reveals within-clonal transcript correlations in yeast populations’, Nature Microbiology, 4(April), pp. 683–692. Available at: 10.1038/s41564-018-0346-9.

Pernice, M.C. et al. (2016) ‘Large variability of bathypelagic microbial eukaryotic communities across the world’s oceans’, ISME Journal, 10(4), pp. 945–958.

Picelli, S. et al. (2014) ‘Full-length RNA-seq from single cells using Smart-seq2’, Nature Protocols, 9(1), pp. 171–181.

Poran, A. et al. (2017) ‘Single-cell RNA sequencing reveals a signature of sexual commitment in malaria parasites’, Nature, 551(7678), pp. 95–99.

Reid, A.J. et al. (2018) ‘Single-cell RNA-seq reveals hidden transcriptional variation in malaria parasites’, eLife, 7, pp. 1–29.

Rognes, T. et al. (2016) ‘VSEARCH: A versatile open source tool for metagenomics’, PeerJ, 2016(10), pp. 1–22.

Rubino, F. et al. (2014) ‘MGkit: Metagenomic Framework For The Study Of Microbial Communities’. Available at: https://bitbucket.org/setsuna80/mgkit.

Saint, M. et al. (2019) ‘Single-cell imaging and RNA sequencing reveal patterns of gene expression heterogeneity during fission yeast growth and adaptation’, Nature Microbiology, 4(3), pp. 480–491.

Simpson, A.G.B., Slamovits, C.H. and Archibald, J.M. (2017) ‘Protist Diversity and Eukaryote Phylogeny’, in J.M. Archibald, A.G.B. Simpson, and C.H. Slamovits (eds) Handbook of the Protists. 2nd edn. Cham, Switzerland: Springer, pp. 1–21.

Smith, D.R. and Keeling, P.J. (2016) ‘Protists and the Wild, Wild West of Gene Expression: New Frontiers, Lawlessness, and Misfits’, Annual Review of Microbiology, 70(1), p. annurev-micro-102215-095448.

Turchinovich, A. et al. (2014) ‘Capture and Amplification by Tailing and Switching (CATS), Anultrasensitive ligation-independent method for generation of DNA libraries for deep sequencing from picogram amounts of DNA and RNA’, RNA Biology, 11(7), pp. 817–828.

Varadi, M. et al. (2022) ‘AlphaFold Protein Structure Database: Massively expanding the structural coverage of protein-sequence space with high-accuracy models’, Nucleic Acids Research, 50(D1), pp. D439–D444.

Vass, M., Székely, A.J., Lindström, E.S. and Langenheder, S. (2020) ‘Using null models to compare bacterial and microeukaryotic metacommunity assembly under shifting environmental conditions’, Scientific Reports, 10(1), pp. 1–13.

Venter, J.C. et al. (2004) ‘Environmental Genome Shotgun Sequencing of the Sargasso Sea’, Science, 304(5667), pp. 66–75.

Walker, G., Dorrell, R.G., Schlacht, A. and Dacks, J.B. (2011) ‘Eukaryotic systematics: a user’s guide for cell biologists and parasitologists’, Parasitology, 138(13), pp. 1638–1663.

Worden, A.Z. et al. (2015) ‘Rethinking the marine carbon cycle: Factoring in the multifarious lifestyles of microbes’, Science, 347(6223).

Xue, Y. et al. (2020) ‘A single-parasite transcriptional atlas of *Toxoplasma gondii* reveals novel control of antigen expression’, eLife, 9, pp. 1–27.

Yilmaz, P. et al. (2014) ‘The SILVA and “all-species Living Tree Project (LTP)” taxonomic frameworks’, Nucleic Acids Research, 42(D), pp. 643–648.

