## Supplementary Material for "Single-cell transcriptomics reveals functional insights into a non-model aquatic phytoflagellate and its metabolically linked bacterial community"

### Supplementary information

#### Supplementary figures

Supplementary Figure 1. Abundance of indigenous bacterial communities in co-culture with Ochromonas triangulata. Time (days) refers to the time elapsed since the last culture passing. Batch B14 was the source of the slow-growing *O. triangulata* cell consortium (sampled at day 12, not shown here), while batch B15 was the source of the fast-growing (sampled at day 2). Shaded areas indicate the concurrent fast and slow growth phases of *O. triangulata*.

Supplementary Figure 2. LysoSensor Blue (LS) staining as a proxy for cell viability in Ochromonas triangulata. Double staining with LS and fluorescein diacetate (FDA), a commonly used reporter for cell viability (MacIntyre and Cullen 2016) shows that both LS and FDA reveal identical populations of living cells. Cells were heat-killed by immersing cultures in a water bath at 55°C during 15 min, then tempered at 18°C during 30 min prior to staining. Stain final concentrations were FDA 1 μg/mL, LS 0.5 mM. (+) Stain added, (–) stain absent. FDA solutions were freshly prepared and used within 30 min from preparation. All scatterplots show blue fluorescence intensity (EX 405, EM 450/45) on the horizontal axis and green fluorescence intensity (EX 488, EM 525/40) on the vertical axis.

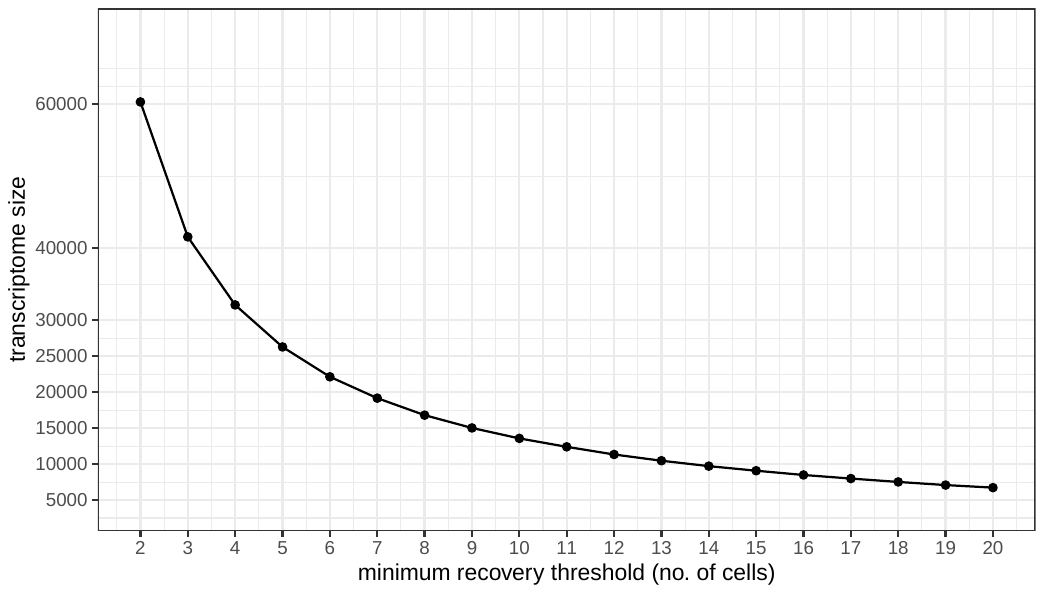

Supplementary Figure 3. Resulting transcriptome size (number of unique transcripts) after successive minimum recovery thresholds (MRTs). The MRT represents the minimum number of cells required to feature a given transcript in order to legitimate the presence of the transcript in the transcriptome. In our analysis, only transcripts recovered at MRT of 2 cells were considered.

Supplementary Figure 4. Relative proportion of annotated and unannotated transcripts. Cluster identity corresponds to postclustering affiliation.

Supplementary Figure 5. Relative proportion of total number of reads that either didn’t map to the assembled transcriptome, were assigned as ribosomal or mapped to either unannotated or annotated transcripts. Cluster identity corresponds to postclustering affiliation.

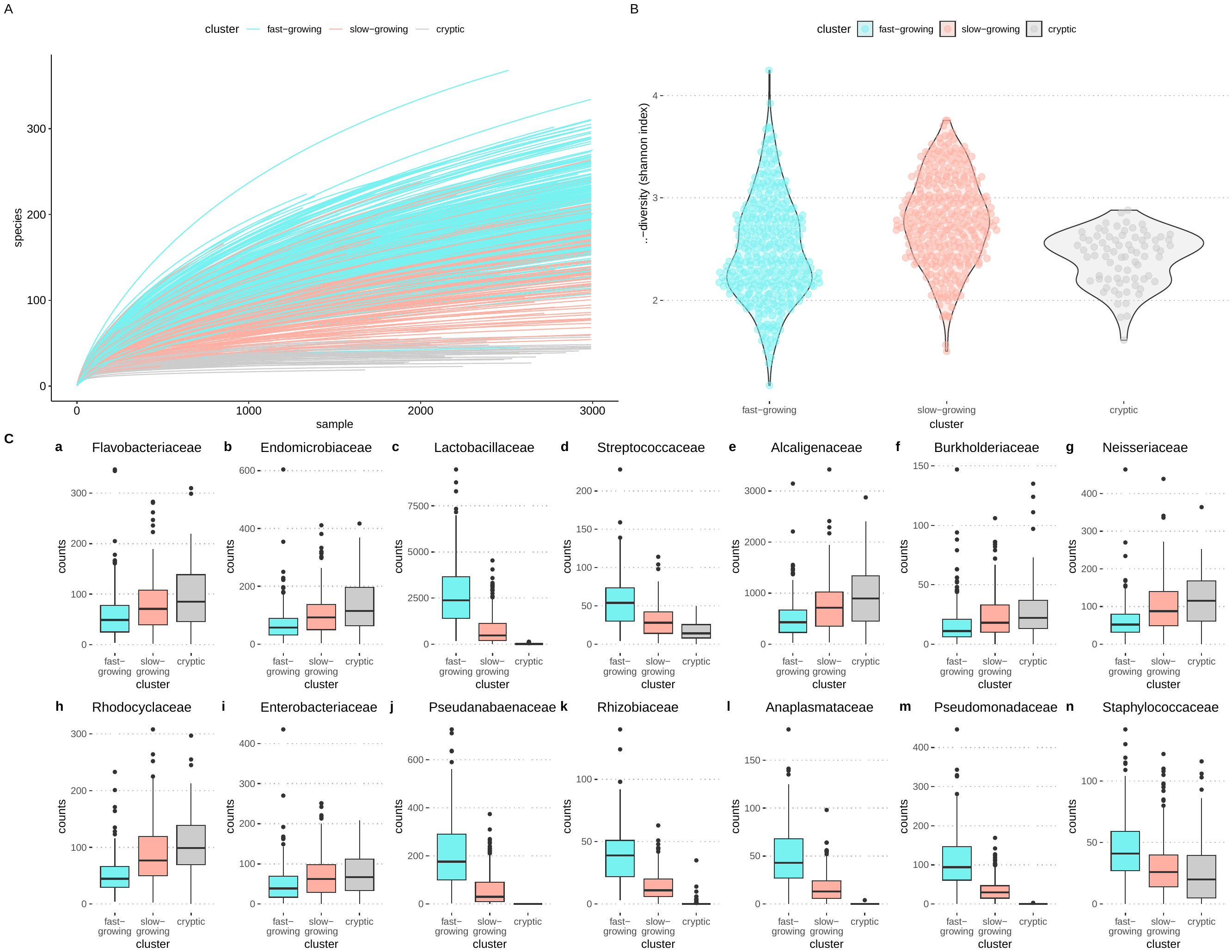

Supplementary Figure 6. Prokaryota associated with the *O. triangulata* cells in culture. (A) Rarefaction curves. (B) Alpha diversity in the three cell clusters as measured using Shannon index. (C) Read counts of the differentially abundant families of prokaryota.

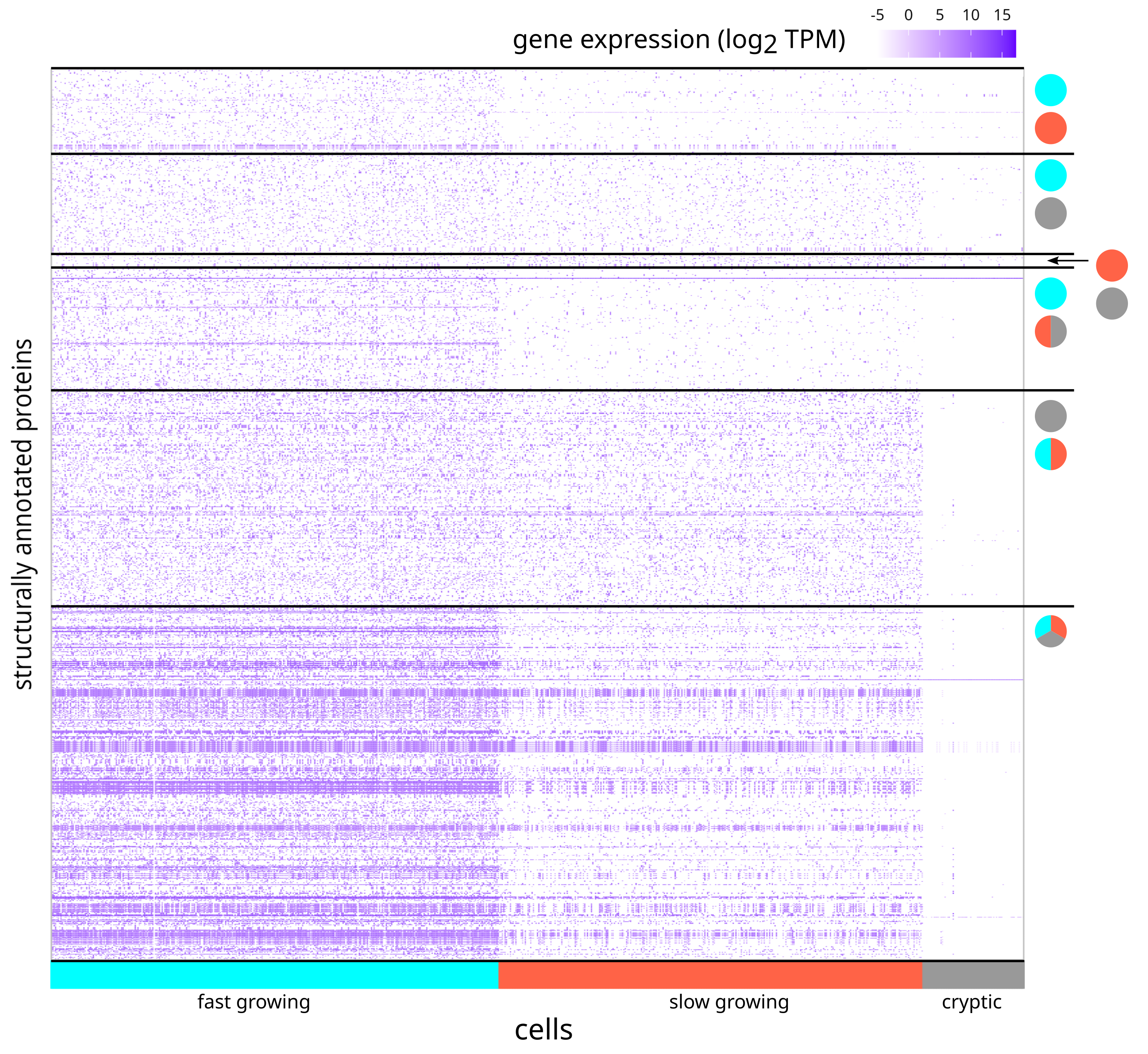
Supplementary Figure 7. Heatmap of structurally-annotated proteins that were differentially expressed between different Ochromonas triangulata growth stages. Coloured rectangles in the horizontal axis indicate cell affiliation to clustering group. Circle pairs indicate pairwise comparisons between stages. Those cases where gene expression in one group is significantly different to that of the other two groups, the joint is represented by a split circle in which the colour of each half encodes the identity of each member. A three coloured circle represents cases in which gene expression differs significantly for any possible pairwise comparison. The colours in circles and rectangles correspond to fast-growing (cyan), slow-growing (red), and cryptic (grey) stages.

#### Supplementary tables

Supplementary Table 1. Size distribution of the proteins in the three clusters of cells. Size refers to no. of amino acids: small (0, 200], medium (200, 400], large (400, 800].

Supplementary Table 2. Quality distribution of the proteins in the three clusters of cells. Quality refers to pLDDT scores: good (0.5, 0.7], high (0.7, 0.9], very high (0.9, 1].

Supplementary Table 3. Significantly enriched Gene Ontology: Biological Processes genesets with the pre-ranked list being ranked based on the wald statistic from the differential expression test between fast-growing and slow-growing clusters.

Supplementary Table 4. Significantly enriched Gene Ontology: Biological Processes genesets with the pre-ranked list being ranked based on the wald statistic from the differential expression test between fast-growing and cryptic clusters.

Supplementary Table 5. Significantly enriched Gene Ontology: Biological Processes genesets with the pre-ranked list being ranked based on the wald statistic from the differential expression test between slow-growing and cryptic clusters.
